# Mitochondrial Dysfunction in Alzheimer’s Disease Is Driven By Excess ER-Calcium Release in Patient-Derived Neurons

**DOI:** 10.1101/2024.10.11.617867

**Authors:** Sarah Mustaly-Kalimi, Wacey Gallegos, Robert A. Marr, Daniel A. Peterson, Israel Sekler, Grace E. Stutzmann

**Author notes:** Corresponding author: Grace E. Stutzmann 3333 Green Bay Rd. North Chicago, IL 60064. **Competing Interest Statement:** The authors have no competing interests.

## Abstract

Tight regulation of mitochondrial Ca^2+^ is essential for neuronal bioenergetics and cellular metabolism. Ca^2+^ transfer from ER-localized ryanodine receptors (RyR) and inositol triphosphate receptors (IP_3_R) to the mitochondria maintains a steady Ca^2+^ source that fuels oxidative phosphorylation and ATP production. In Alzheimer’s disease (AD), RyR-evoked Ca^2+^ release is markedly increased, contributing to synaptic deficits, protein mishandling, and memory impairment. Here, we demonstrate that dysregulated RyR-Ca^2+^ release directly compromises mitochondrial activity and is an early contributor to AD cellular pathology. We measured an array of mitochondrial functions using fluorescent biosensors and optical imaging in RyR2-expressing HEK cells and iPSC-derived neurons from familial AD and nonAD patients. In neurons from AD patients, resting mitochondrial Ca^2+^ levels were elevated alongside increased free radical production and higher caspase-3 activity relative to nonAD neurons. RyR-evoked Ca^2+^ release further potentiated pathogenic mitochondrial responses in AD neurons, with increased Ca^2+^ uptake and exaggerated membrane depolarization. Additionally, clearance of damaged mitochondria was impaired in AD neurons, demonstrating consequences from dysfunctional lysosomes. Notably, impairments to mitochondria in AD neurons were largely prevented with the RyR negative allosteric modulator, Ryanodex. These findings highlight how excess RyR-Ca^2+^ release broadly contributes to early cellular pathology in AD which includes a cascade of ER, lysosomal, and mitochondrial deficits culminating in neuronal decline and degeneration. Additionally, pharmacological suppression of RyR-Ca^2+^ release preserves mitochondrial, ER and lysosomal function, thus providing a novel and effective therapeutic.

**Significance Statement:** Mitochondrial dysfunction plays a central role in the cellular pathogenesis of Alzheimer’s disease, yet the upstream mechanisms driving these deficits remain largely unknown. Here, in human neurons derived from Alzheimer’s patients, we reveal how early Ca^2+^ mishandling through the ER-localized ryanodine receptor (RyR) is sensed by the mitochondria and triggers a pathogenic metabolic cascade. Pharmacologically restoring Ca^2+^ homeostasis reversed these deficits, including the increased free radicals and defective mitophagy, and restored the nonAD phenotype. The findings reveal the early neuronal signaling mechanisms that affect ER Ca^2+^ handling, proteolysis, and mitochondrial activity, highlighting the potential role of the dysregulated RyR as a therapeutic target for AD using clinically relevant model systems.

## Introduction

Mitochondria are vital organelles that regulate cellular bioenergetics and support the high metabolic demands of neurons. An equally essential role of this organelle is maintaining homeostatic regulation of free Ca^2+^ levels within the cytosol. Along with the ER and lysosomes, mitochondria store and buffer intracellular Ca^2+^ to shape Ca^2+^ signals and support a range of physiological functions, including synaptic transmission and plasticity (1–3), gene expression (4, 5), and ATP production (6–8).

Ca^2+^ is driven into the mitochondria by its highly hyperpolarized membrane potential (ΔΨ_m_; -150 to -180mV), passing through the voltage-dependent anion channel (VDAC) in the outer mitochondrial membrane (OMM) and then into the matrix through the selective inner mitochondrial membrane (IMM)-localized mitochondrial Ca^2+^ uniporter (MCU) (9, 10). At rest, the Ca^2+^ concentration within the matrix ([Ca^2+^]_m_) is approximately 100-200nM, with levels rising to 1-10μM during Ca^2+^ spikes (3, 10–12). While the mitochondria can tolerate increases in [Ca^2+^] of several orders of magnitude during bursts of neuronal activity, prolonged mitochondrial Ca^2+^ influx can feed into a pathological cascade of events, including depolarized membrane potential, increased reactive oxygen species (ROS) production, and release of apoptotic initiators such as cytochrome c and caspase activation (2, 13–19).

Elevated ryanodine receptor (RyR) mediated Ca^2+^ release from ER stores is a well-documented early pathogenic mechanism in AD observed across various mouse and human models (20–24). The excessive Ca^2+^ release contributes to deficits in synaptic structure and function (25), impaired lysosome proteolytic capacity (26–28), hyperphosphorylation of tau and Aβ_42_ aggregation, and cognitive decline (23, 27, 29, 30). The tight coupling between the ER and mitochondria renders the mitochondria especially vulnerable to elevated RyR-Ca^2+^ release. Furthermore, the redox-sensitive RyR2 isoform is upregulated in AD-vulnerable brain regions (22, 31, 32) and becomes sensitized and leaky through post-translational modifications such as oxidation and phosphorylation, thus propagating a feed-forward cycle of aberrant ER-Ca^2+^ release and mitochondrial Ca^2+^ overload.

Despite the increasing focus on the role of mitochondrial dysfunction and oxidative stress in AD (33–35), the upstream pathogenic mechanisms driving mitochondrial stress remain elusive. We propose that altered ER-Ca^2+^ signaling drives mitochondrial-mediated cellular pathophysiology and Ca^2+^ dyshomeostasis. In the present study, we demonstrate a pathogenic inter-organelle coupling dynamic and show how increased RyR-Ca^2+^ release causes mitochondrial dysfunction through increased Ca^2+^ load, leading to a depolarized mitochondrial membrane potential, increased superoxide production, elevated caspase-3 activity and defective mitophagic clearance in iPSC-derived neurons from AD patients and age/sex-matched controls. Importantly, we further demonstrate the therapeutic potential of normalizing RyR-Ca^2+^ signaling, thereby identifying an upstream mechanism-based strategy for preventing AD progression.

## Results

Given the known ER-Ca^2+^ handling alterations in AD, we first sought to determine if the mitochondria detect this excess Ca^2+^ and, if so, investigate its effects on mitochondrial functions contributing to AD-related deficits. Furthermore, we wish to validate therapeutic strategies that target upstream cellular mechanisms driving cognitive deficits, neuronal toxicity, and impaired proteostasis in AD.

### Elevated RyR-Ca^2+^ Release Increases Mitochondrial Ca^2+^ Influx in Model Cells and Human iPSC-derived Neurons from AD Patients

Here, we investigated if RyR-Ca^2+^ release is a source of mitochondrial Ca^2+^ influx and an initiating factor of mitochondrial dysfunction. To accomplish this, we measured mitochondrial matrix Ca^2+^ levels using AAV9-hSyn-2mt-Mito-GCaMP6m, a genetically encoded Ca^2+^ indicator tagged to the inner mitochondrial membrane. In RyR2-expressing HEK (RyR2 HEK) cells, caffeine-evoked RyR-Ca^2+^ release was followed by increased mitochondrial Ca^2+^ levels and was inhibited by Ryanodex (10μM; 1hr), a negative allosteric RyR modulator (Fig. 1B&D: n=9 control coverslips; n=8 Ryanodex coverslips; t_(15)_ = 4.14; p<0.0001). Ryanodex treatment did not alter resting mitochondrial Ca^2+^ levels (Fig.1A&C: n=9 control; n=8 Ryanodex; t_(15)_=2.11; p=0.05). Mitochondrial Ca^2+^ influx rate was reduced after Ryanodex treatment (Fig.1E: n=9 control; n=8 Ryanodex; t_(15)_ =2.43; p=0.03), but did not alter mitochondrial Ca^2+^ efflux rate (Fig.1F: n=9 control; n=8 Ryanodex; t_(15)_ =0.73; p=0.48). These data demonstrate that mitochondrial Ca^2+^ influx is tightly coupled to RyR-Ca^2+^ signaling with a rapid Ca^2+^ transfer between the ER and mitochondria.

**Figure 1.**
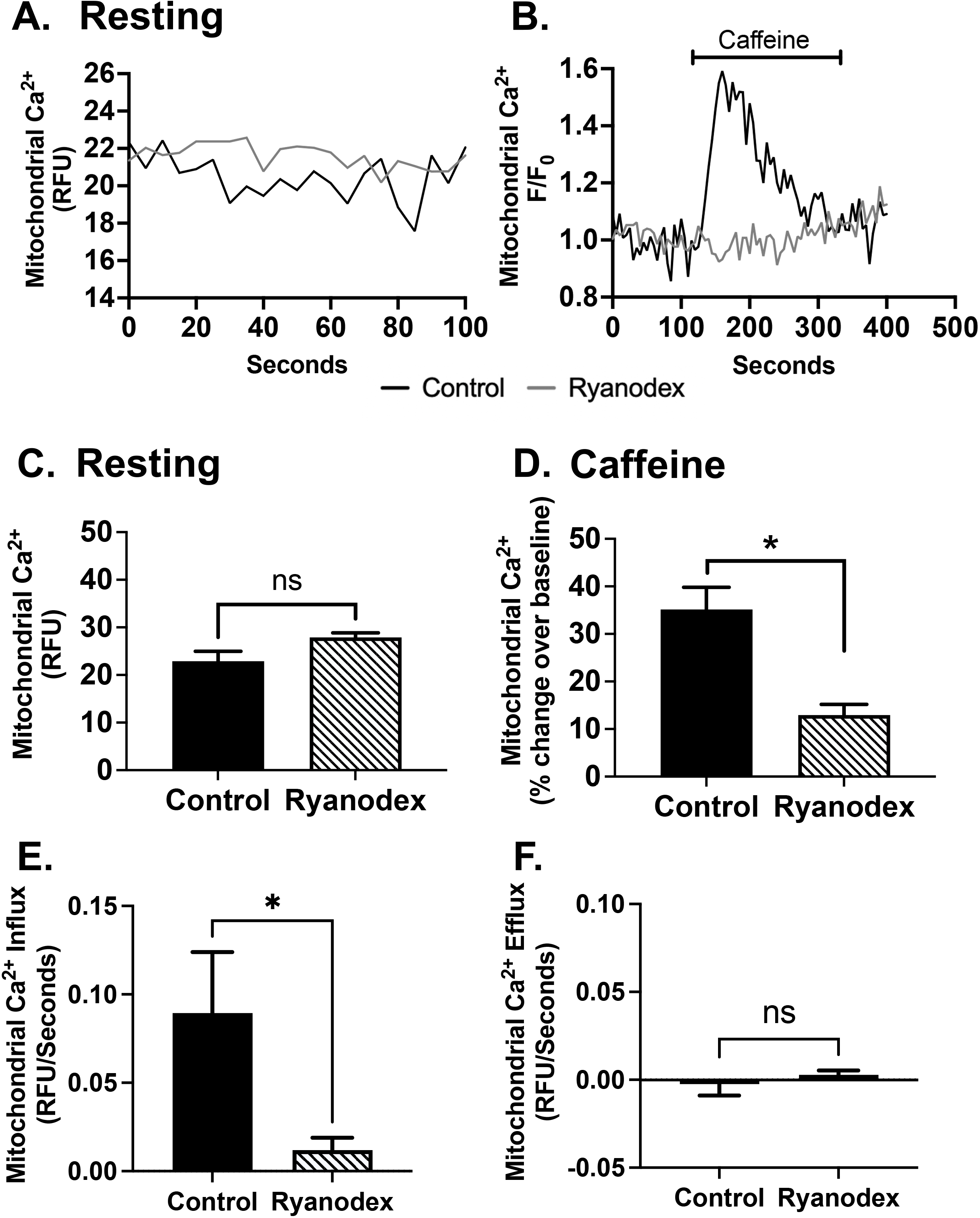
Ryanodine Receptor (RyR2)-evoked Ca^2+^ release drives mitochondrial Ca^2+^ influx in model cells. Cells were incubated with crude lysate of AAV9-hSyn-2mt-Mito-GCaMP6m to measure mitochondrial Ca^2+^ levels in HEK293T cells expressing hippocampal-relevant RyR2 cells. Representative traces of mitochondrial Ca^2+^ levels during (A) rest (relative fluorescence) and (B) time-course recordings (fluorescence normalized to rest). (C) At rest, no significant difference is seen in mitochondrial Ca^2+^ levels between treatment groups. (D) Normalizing RyR2-Ca^2+^ release with Ryanodex treatment (10μM; 1 hour) attenuated mitochondrial Ca^2+^ levels upon evoked RyR-Ca^2+^ release. (E) Ryanodex treatment reduced mitochondrial influx relative to control, but (F) mitochondrial Ca^2+^ efflux was not significantly changed. (control: n=9, Ryanodex: n=8 coverslips/treatment) *p<0.05; error bars describe SEM. Relative fluorescence unit (RFU)

To incorporate translationally relevant models of AD, we next utilized human iPSC-derived neurons (HiNs) generated from AD patients and age- and sex-matched healthy controls (nonAD). Using the genetically encoded indicator described above, mitochondrial Ca^2+^ levels were measured at resting/baseline (Fig. 2A&C) and in response to caffeine-evoked (10mM) RyR-Ca^2+^ release (Fig. 2B&D). At baseline, the steady-state Ca^2+^ load was significantly higher in AD compared to nonAD HiN by 46.72 ± 1.06%, with a significant reduction in resting Ca^2+^ levels after normalizing RyR-Ca^2+^ release with Ryanodex (Fig. 1C: n=72 nonAD neurons; n=82 AD neurons, F_(3,304)_=18.95, p<0.0001). Notably, Ryanodex treatment in AD HiNs did not fully restore mitochondrial Ca^2+^ to nonAD levels, suggesting that additional sources or mechanisms may contribute to elevated resting Ca^2+^ levels, such as lysosomal Ca^2+^ stores or defective clearance through the NCLX extrusion channels. With evoked RyR-Ca^2+^ release, mitochondrial matrix Ca^2+^ levels further increased in the AD HiNs compared to nonAD HiN (Fig. 2D: n=72 nonAD neurons, n=82 AD neurons, F_(3,304)_=13.53, p<0.0001). Ryanodex treatments restored mitochondrial Ca^2+^ uptake in the AD HiN to nonAD levels, with no notable effect in the nonAD HiNs. Mitochondrial Ca^2+^ influx rate was increased in AD HiN, which was attenuated with Ryanodex treatment and comparable to nonAD HiN rates (Fig. 2E: n=72 nonAD neurons, n=82 AD neurons, F_(3,304)_ = 5.96, p=0.0006) Mitochondrial Ca^2+^ efflux rate was greater in AD HiN, regardless of Ryanodex treatment (Fig. 2F: n=72 nonAD neurons, n=82 AD neurons, F_(3,304)_ = 7.27, p<0.0001).

**Figure 2:**
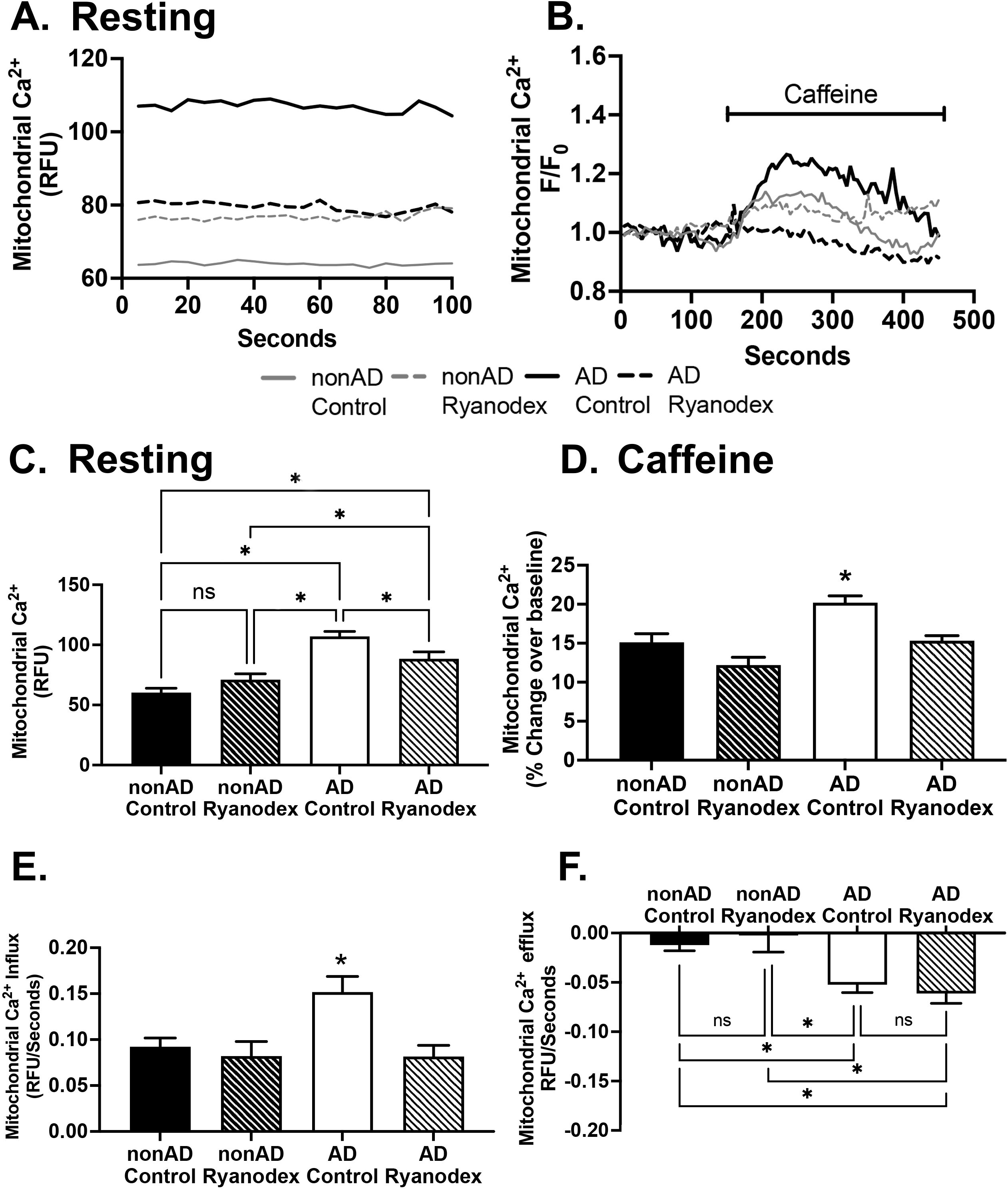
Ryanodine Receptor (RyR)-evoked Ca^2+^ release drives mitochondrial Ca^2+^ influx in human neurons. Cells were incubated with crude lysate of AAV9-hSyn-2mt-Mito-GCaMP6m to measure mitochondrial Ca^2+^ levels in human-induced neurons from nonAD and AD patients. Representative traces of mitochondrial Ca^2+^ levels during (A) rest (relative fluorescence) and (B) time-course recordings (fluorescence normalized to rest). (C) Resting mitochondrial Ca^2+^ levels were increased in control AD HiNs with partial rescue of mitochondrial Ca^2+^ levels to nonAD in Ryanodex-treated AD HiNs. (D) Caffeine-evoked RyR-Ca^2+^ release increased mitochondrial Ca^2+^ levels in AD HiNs, which was attenuated to nonAD levels with Ryanodex treatments (10μM; 1 hour). (E) Mitochondrial Ca^2+^ influx was elevated in AD HiN, which was attenuated with Ryanodex treatment. (F) Mitochondrial Ca^2+^ efflux was greater in AD neurons compared to nonAD, regardless of Ryanodex treatment. (nonAD: n=72, AD: n=82 neurons/treatment) *p<0.05; error bars describe SEM. Relative fluorescence unit (RFU)

Here, we demonstrate a close coupling between RyR-Ca^2+^ release and mitochondrial Ca^2+^ uptake in model cells and HiNs, with exaggerated responses in the AD HiNs. Notably, at rest, Ca^2+^ levels within the mitochondrial matrix were increased in the AD HiNs, implicating a steady-state Ca^2+^ leak from the ER and possibly other organelles in AD neurons.

### RyR-Ca^2+^ Release Depolarizes Mitochondrial Membrane Potential in Model Cells and Human Neurons

The strong mitochondrial electrochemical gradient (ΔΨ_m_; -150mV to -180mV) drives Ca^2+^ into the matrix and serves as an energy store for oxidative phosphorylation (36). Activity-dependent transient increases in mitochondrial Ca^2+^ will temporarily depolarize the ΔΨ_m_, with Ca^2+^ efflux through Na^+^/Ca^2+^/Li^+^ exchanger (NCLX) restoring membrane potential. However, prolonged or excessive Ca^2+^ influx into the mitochondrial matrix may cause maladaptive ΔΨ_m_ depolarization. Since mitochondrial Ca^2+^ levels are elevated in the AD HiNs at baseline and RyR-activated conditions, we sought to determine the impact of elevated RyR-Ca^2+^ signaling on mitochondrial ΔΨ_m_ in RyR2 HEK cells and HiNs.

Baseline (resting) ΔΨ_m_ was measured in RyR HEK cells and human neurons using tetramethylrhodamine methyl ester in quenching mode (TMRM; 50nM, 1hr), which is an optimal sensor for steady-state readouts (37). When normalized to the CCCP-generated maximal fluorescence, resting ΔΨ_m_ in RyR2 HEK cells did not significantly change in response to Ryanodex treatment indicating little or no direct effect of the drug on ΔΨ_m_ (**Fig. 3A-C**; (t_(8)_=1.80; p=0.42; control: n=5 coverslips; Ryanodex: n=3 coverslips). Caffeine-mediated RyR activation did generate a significant depolarization as measured by TMRM (t_(8)_=4.43; p<0.05) although this was repeated using Rhod123 which is a preferable method to quantify dynamic ΔΨ_m_ changes (37) (see below). Similarly, in the human neurons, there were no significant differences in the TMRM-measured resting ΔΨ_m_ among groups when normalized to the CCCP maximum response (Fig. 3D-F; F_(3,440)_=1.79; p=0.15; Controls: n=112 nonAD neurons, n=102 AD neurons; Ryanodex: n=127 nonAD neurons & n=99 AD neurons), and activating RyR with caffeine resulted in an greater membrane depolarization in the AD HiN which was normalized with Ryanodex (p<0.05; data not shown; see below for Rhod123 analysis). As a positive control, CCCP-mediated uncoupling showed maximal depolarization in both human neurons and RyR2 HEK cells with no significant differences observed among experimental conditions or groups (Fig 3C&F; F_(3,440)_=1.54; p=0.20).

**Figure 3.**
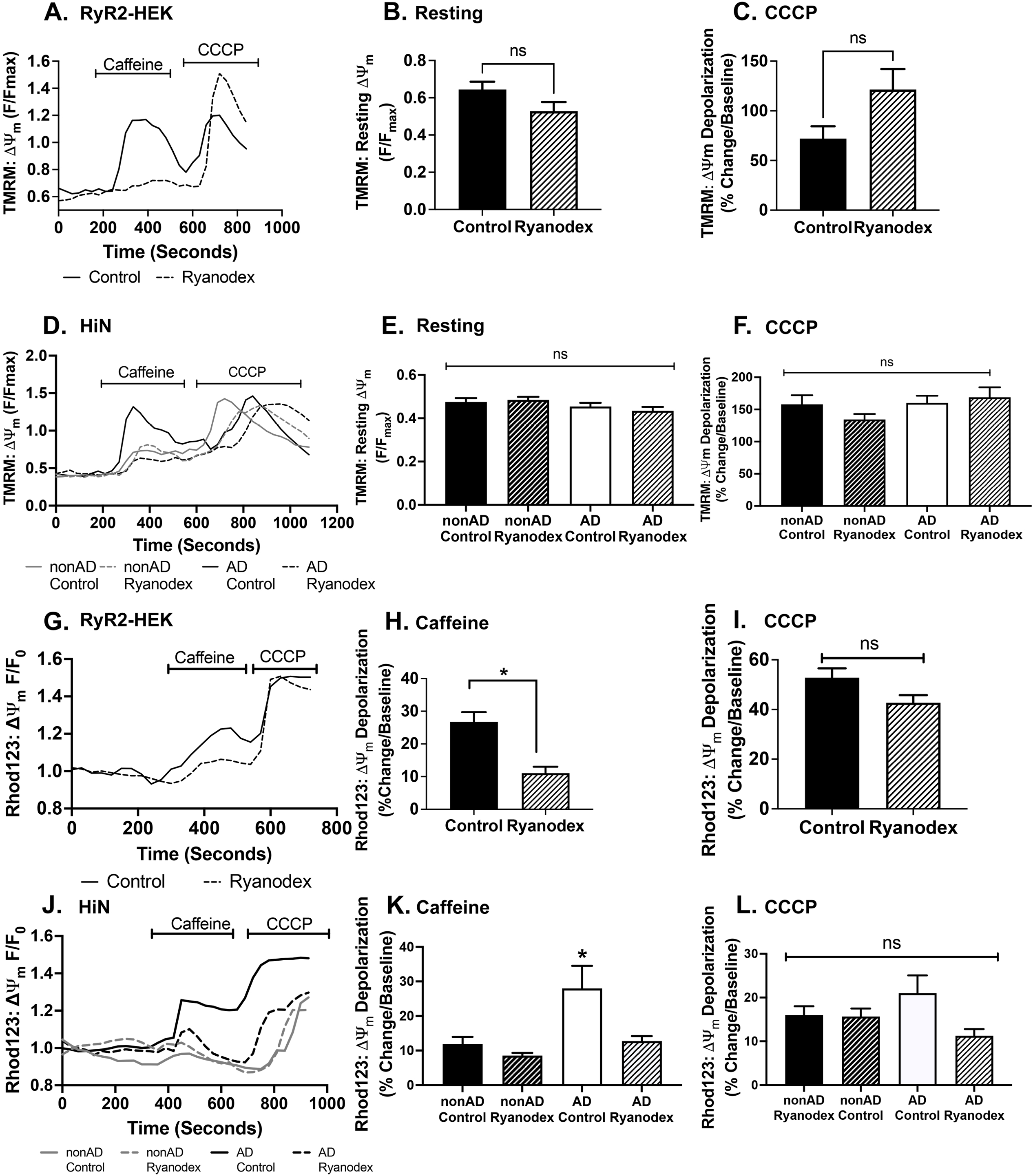
Ryanodine Receptor (RyR)-evoked Ca^2+^ release depolarizes mitochondrial membrane potential in model cells and human neurons Depolarization of the mitochondrial membrane potential (ΔΨ_m_) was both ([10μM] Rhod123 or [50nM] TMRM). (A,D,G, J) Representative traces of time-course recordings measuring (ΔΨ_m_) in response to RyR-Ca^2+^ release (10mM caffeine) and 5μM CCCP (A: RyR2 HEK with TMRM; D HiNs with TMRM, G: HEK-RyR2 cells with Rhod123, J: HiNs with Rhod123). (B) When measuring with TMRM, there was no significant difference in Ψ_m_ at rest in both RyR2 HEK (controls: n=5; Ryanodex: n=3; n=coverslips) (B) and HiN (Controls: n=112 nonAD neurons, n=102 AD neurons; Ryanodex: n=127 nonAD neurons & n=99 AD neurons) (E) after normalizing to CCCP-generated maximum fluorescence. When measuring with Rhod123, fluorescence was normalized to baseline to determine membrane potential changes from resting. (H) In RyR2 HEK cells after normalizing RyR-Ca^2+^ release with Ryanodex attenuated mitochondrial membrane depolarization in response to caffeine-evoked RyR-Ca^2+^ release(control n=8, Ryanodex n=8 coverslips). (K) In HiN, In human neurons, caffeine-evoked RyR-Ca^2+^ release triggered a significant depolarization of the Ψ_m_ in AD control HiNs compared to all other groups, with Ryanodex treatments reducing depolarization to nonAD levels (nonAD control n=86, nonAD Ryanodex n=91, AD control n=34, AD Ryanodex n=52 neurons). There was no significant changes in fluorescent signaling in all groups after CCCP collapse of ΔΨ_m_ in both (I) RyR HEK and (L) HiN. *p<0.05; errors bars describe SEM

The cationic lipophilic dye rhodamine123 (Rhod123: 10μM, 30min) is a preferred method for capturing dynamic changes to the ΔΨ_m_ (37), such as in response to RyR activation by caffeine. Here, evoked responses are calculated relative to their respective resting baseline. In RyR2 HEK cells, caffeine-mediated RyR2 activation significantly depolarized the mitochondrial membrane potential, and this effect was was blunted by Ryanodex (10mM; 1hr) (Fig. 3G-I: n=8 coverslips; t_(14)_ = 4.37; p=0.0006). CCCP-mediated uncoupling resulted in maximal membrane depolarization and was not affected by Ryanodex (Fig. 3I: t_(14)_=2.10; p>0.05). In the human neurons, caffeine-evoked RyR-Ca^2+^ release resulted in greater ΔΨ_m_ depolarization in AD neurons compared to nonAD neurons, with Ryanodex treatment restoring membrane depolarization in AD conditions to nonAD level; Ryanodex did not affect nonAD neurons membrane potential suggesting no detectable degree of RyR leak under healthy conditions (Fig. 3J&K F_(3,259)_=9.21, p<0.0001; Controls: n=86 nonAD neurons, n=34 AD neurons; Ryanodex: n=91 nonAD neurons, 52 AD neurons). CCCP-induced uncoupling showed maximal depolarization in human neurons, with no differences observed among experimental conditions or groups (Fig. 3L: F_(3,255)_=2.00, p=0.11). Together, these data show that the RyR-Ca^2+^ dysregulation associated with AD provides excessive Ca^2+^ to the mitochondria and disproportionally depolarizes the ΔΨ_m_. Restoration to a physiological ΔΨ_m_ with Ryanodex highlights the key role of ER-Ca^2+^ dyshomeostasis in disrupting mitochondrial function in AD.

### Elevated Superoxide Production in AD Human Neurons is Normalized by Attenuating RyR-Ca^2+^ Release

Excess free radicals are detrimental to cellular activity due to their ability to oxidize fatty acids, protein complexes, and nucleotides, thus accelerating disease progression (38). To determine if the increased mitochondrial Ca^2+^ uptake leads to increased oxidative free radicals in AD, superoxides were measured in AD and nonAD HiN using the fluorogenic dye MitoSOX (10μM). Under basal conditions, superoxide production was significantly increased in the AD HiN relative to nonAD, and Ryanodex pre-treatment (10μm; 1hr) reduced superoxide production in the AD neurons to nonAD levels, with no significant effects in the nonAD neurons (Fig 4A&B: F_(3,28)_=10.50. P<0.0001, n=8 coverslips). To control for possible differences in fluorescence measurements due to cell density, cells were counted for each experimental condition, with no significant difference between the groups (Supplemental Figure 1A: average cell count 42.0 ± 16.5 cells per well F_(3,29)_=1.58, p=0.21).

**Figure 4.**
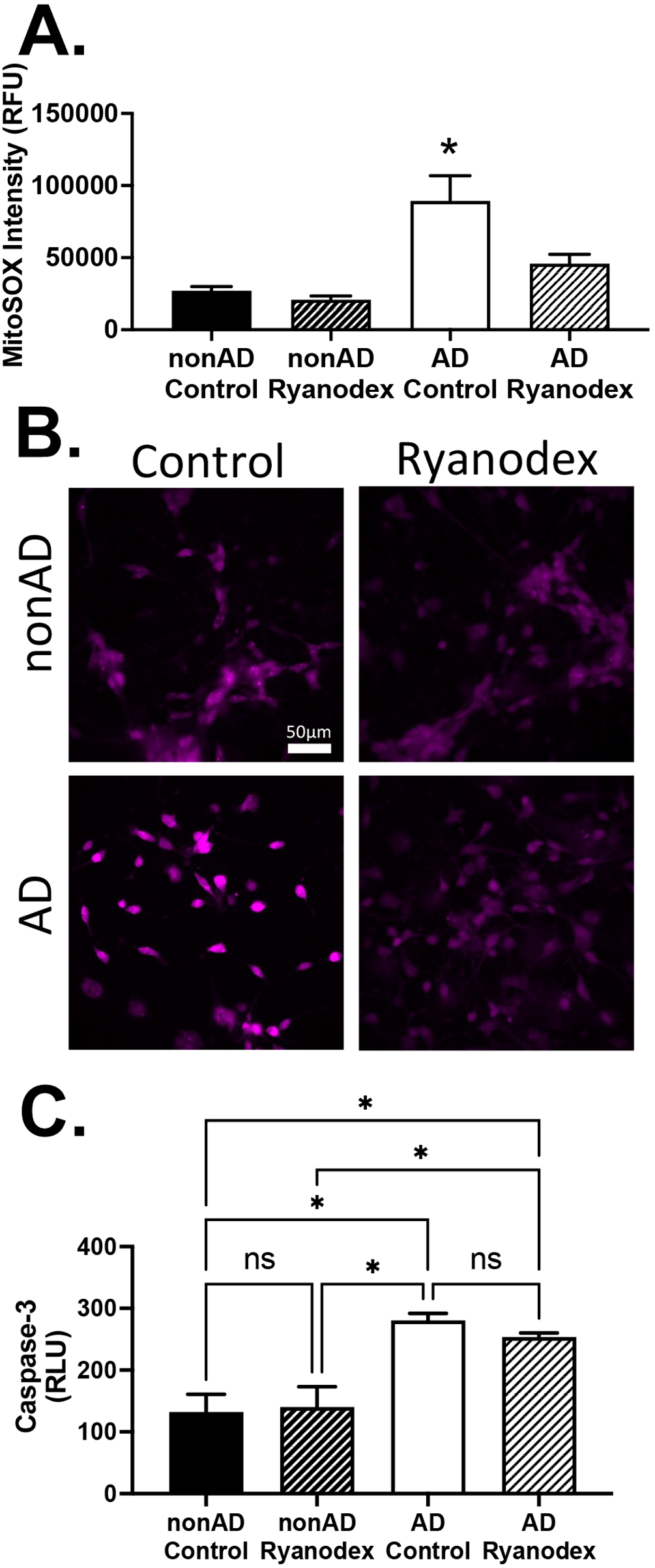
Increased superoxide production and caspase-3 activity in human AD neurons (A) Representative images of mitochondrial superoxide production measured by MitoSOX dye in nonAD and AD HiNs treated with and without Ryanodex (10μM; 1 hour). (B) Quantification of fluorescent intensity shows increased mitochondrial superoxide production in AD HiNs, which was normalized to nonAD levels with Ryanodex treatments (n=8 coverslip/treatment). (C) Caspase-3 activity is significantly higher in AD HiNs compared to nonAD HiNs, and Ryanodex (10μM 6 hours) treatment did not significantly impact caspase-3 activity in AD neurons (n=12 wells/treatment). *p<0.05; error bars describe SEM. Relative fluorescence unit (RFU) Relative luminescence unit (RLU)

Here, we demonstrate a direct coupling between aberrant RyR-Ca^2+^ leak and increased mitochondrial superoxide production in AD neurons. As this was reversed with Ryanodex, only in AD neurons, this suggests this RyR-Ca^2+^ signaling defect is not present under normal physiological resting conditions and is a unique feature of AD. Furthermore, it has previously been shown that oxidation of the RyR2 channel at free cysteine residues alters its secondary protein structure and results in increased Ca^2+^ release (22, 39–43), thus establishing a self-propagating feed-forward loop between excess ER Ca^2+^ release, increased mitochondrial free radical generation, oxidized RyR and excess ER Ca^2+^ leak.

### Increased Caspase-3 Activity in AD Neurons

Mitochondrial Ca^2+^ overload can initiate apoptotic cell death pathways via opening of the mitochondrial permeability transition pore (mPTP), which releases cytochrome c and activates caspases. To determine if RyR-mediated increases in mitochondrial Ca^2+^ feed into this apoptotic pathway, we measured caspase-3 activity in AD and nonAD neurons using the Caspase-Glo 3/7 assay system (Promega). Consistent with the increase in resting matrix Ca^2+^ levels, caspase-3 activity was significantly greater in AD HiNs compared to nonAD HiNs (Fig. 4C: F_(3,44)_=11.9, p<0.0001). Pre-treatment with Ryanodex (10μM; 6hr) did not significantly reduce the elevated caspase-3 activity in the AD HiNs and had no effect on nonAD HiNs (p>0.05).

These findings suggest that AD HiNs are more susceptible to apoptosis due to increased caspase-3 activity, likely resulting from prolonged mitochondrial stress driven by chronic dysregulated RyR-Ca^2+^ release. In this case, the early disruptions in mitochondrial functions could drive the sustained activation of caspase-3. Thus, Ryanodex exposure would likely have minimal effects on caspase-3 activity within the 6-hour testing window conducted in these studies.

### AD-Associated Increases in ER-Ca^2+^ Release Disrupt Autophagic-Mediated Clearance of Mitochondria (Mitophagy)

Defective mitochondria are removed through mitophagy, a process in which autophagosomes target and engulf the damaged organelle, then fuse to a lysosome to degrade its contents. Disruptions in mitophagy can result from impaired lysosomal digestive capacity, which has been previously demonstrated in AD neurons (27, 44, 45), leading to the accumulation of dysfunctional mitochondria, increased ROS production, impaired mitochondrial respiration, and decreased ATP generation (46–49). In these experiments, we tested whether the disruption to mitochondria function and integrity in response to increased RyR-Ca^2+^ release would increase mitophagy as a protective mechanism, or would mitophagy be impaired due to known defects in lysosomal function.

To measure mitophagic turnover, HiNs were incubated with a mitophagy detection dye (Mitophagy Dye Dojindo 0.1μM) for 30 minutes. We found a significant decrease in mitophagy in AD HiNs compared to nonAD HiNs, with Ryanodex partially resolving the mitophagy deficits in AD HiNs and having no effect on the nonAD neurons (Fig. 5A&B: F_(3,14)_=8.32, p=0.002, n=6). Bafilomycin was used to block the vATPase proton pump on the lysosomes to examine the role of lysosomal acidification in the HiN mitophagy process. Here, mitophagy was largely and equally blocked in both the AD and nonAD neurons, resulting in the conversion of nonAD neurons to the AD phenotype. To determine if the observed reduction in mitophagy was caused by a reduced lysosome pool, lysosomal density was measured using Lysotracker (1μM; 30 mins). No differences in lysosome density were detected between AD and nonAD groups (Fig. 5C: F_(3,14)_=0.89 p = 0.47). Additionally, to control for possible differences in fluorescence measurements due to cell density, cell counts per well were not significantly different amongst treatment groups (Supplemental Figure 1B: average cell count 26.5 ± 4.0 cells per well, (F_(5,21)_=0.48, p = 0.79).

**Figure 5.**
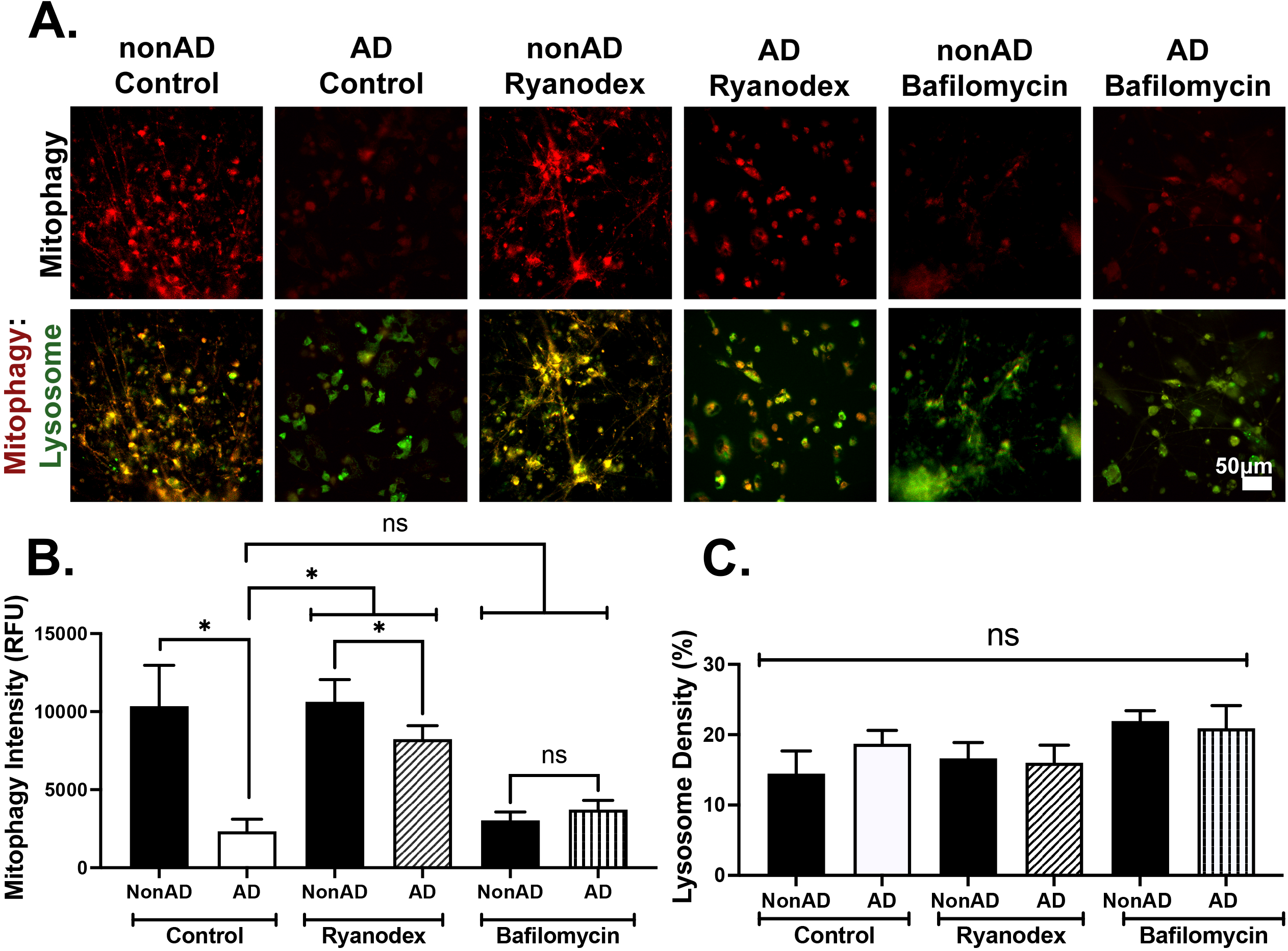
Defective mitophagy is partially rescued in AD neurons after normalizing intracellular Ca^2+^ levels with Ryanodex treatment. (A) Representative images of mitophagy and lysosome density, measured with mitophagy dye and lysosome dye, in nonAD and AD HiNs with and without Ryanodex (10μM; 6 hours) or bafilomycin (inhibitor of vATPase, and positive control for mitophagic inhibition (25nM; 4 hours). (B) Quantification of fluorescent intensity shows decreased mitophagic vesicles in AD HiNs, which is partially rescued after Ryanodex treatment. (C) Quantification of fluorescent intensity of lysosomal density reveals no significant difference in AD HiNs and nonAD HiNs in all treatment groups. (n=6 coverslips/treatment) *p<0.05; error bars describe SEM. Relative fluorescence unit (RFU).

**Figure 6:**
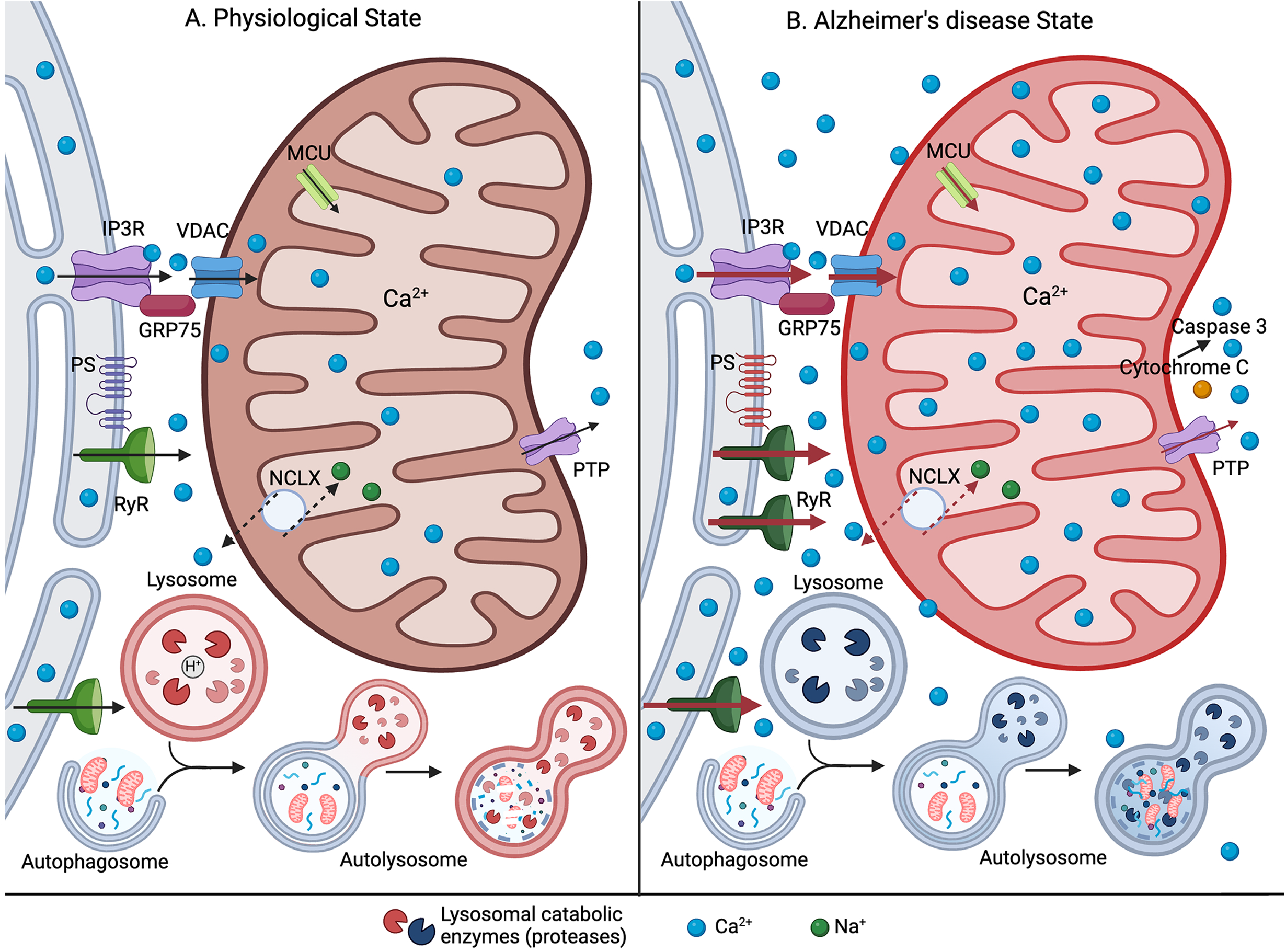
Elevated ER-Ca^2+^ release disrupts mitochondrial calcium handling and bioenergetics in early stages of Alzheimer’s disease. Increased RyR-Ca^2+^ signaling leads to excess Ca^2+^ sequestering into the mitochondria. This increased influx of Ca^2+^ depolarizes the mitochondrial membrane potential. Sustained ionic stress on the mitochondria leads to increased superoxide production and release of apoptotic initiators, like cytochrome c and caspase-3. Additionally, elevated cytosolic Ca^2+^ concentration disrupts the mitophagic clearance of mitochondrial proteins, furthering neuronal stress.

These findings demonstrate that increased RyR-Ca^2+^ release selectively disrupts mitophagy in AD HiNs, and this is largely mediated through impaired lysosomal degradative capacity, as bafilomycin did not further increase mitophagy in the AD neurons, and converted the nonAD neurons to the AD phenotype.

## Discussion

This study demonstrates how aberrant inter-organelle Ca^2+^ signaling contributes to early AD pathophysiology in human neurons. Ca^2+^ dyshomeostasis in AD is a well-established mechanism in various model systems and clinical samples (1, 23, 50–54) and contributes to all major features of the disease, including synaptic loss (55, 56), amyloid and tau aggregation (27, 44, 57), and cell death (58, 59). There has been heightened interest in the role of mitochondrial dysfunction in AD, which underlies hypometabolism (60, 61), increased oxidative free radical generation (16, 62, 63), and synaptic plasticity deficits (64–66). While the consensus is that these deficits play a major role in AD progression, the upstream mechanisms leading to compromised mitochondrial function have remained largely unexplored. In this study, we address this important missing link and provide evidence that excessive RyR-Ca^2+^ release leads to increased mitochondrial Ca^2+^ content, which disrupts a cascade of mitochondrial processes in AD HiNs. This includes increased Ca^2+^ load within the mitochondrial matrix, depolarized mitochondrial membrane potential, increased superoxide production, and upregulated caspase-3 activity, all occurring under resting conditions and further augmented in response to evoked ER-Ca^2+^ release. The interaction with other faulty organelles in AD further accelerates the pathogenic cycle, as seen by a reduction in mitophagy and clearance of damaged mitochondria. Notably, pharmacological suppression of RyR-Ca^2+^ signaling with Ryanodex treatment rescued these deficits in AD human neurons and supports the hypothesis that AD-associated increased RyR-Ca^2+^ signaling is an upstream pathological driver for mitochondrial dysfunction and oxidative stress in AD.

### ER – Mitochondrial Calcium exchange dynamics in AD

The role of the RyR in regulating mitochondrial Ca^2+^ is a focal point in understanding early AD mechanisms, as RyR2 upregulation (31, 67), exaggerated evoked ER-Ca^2+^ release (23, 68–71), and post-translational modifications creating a leaky RyR2 channel (22) are all documented mechanisms associated with AD. The impact of dysregulated RyR-Ca^2+^ signaling is observed in the AD HiNs, with elevated resting mitochondrial Ca^2+^ levels and increased Ca^2+^ uptake in response to caffeine-evoked RyR-Ca^2+^ release compared to nonAD neurons. Using the RyR negative allosteric modulator Ryanodex, elevated RyR-Ca^2+^ signaling is normalized (23) and reduces resting mitochondrial Ca^2+^ levels in AD HiNs (Fig. 2C) with no effect on basal mitochondrial Ca^2+^ levels in nonAD HiNs (Fig. 2C) RyR2 HEK cells (Fig. 1C). We propose that AD-associated RyR-Ca^2+^ leak is the primary mechanism contributing to the elevated resting Ca^2+^ levels observed in AD neurons. Despite reducing resting mitochondrial Ca^2+^ levels following Ryanodex treatments, the mitochondrial Ca^2+^ load within AD neurons remains elevated compared to nonAD conditions (Fig. 2C-D). This sustained elevation of resting Ca^2+^ levels in Ryanodex-treated AD neurons suggests deficits in mitochondrial Ca^2+^ efflux mechanisms, such as through the Na^+^/Ca^2+^ exchanger (NCLX). Impaired NCLX function leads to increased mitochondrial Ca^2+^ levels, accelerated cognitive decline, increased amyloidosis, and tau pathology, thus contributing to AD pathology (2, 17, 72, 73). Upregulated MCU activity may also facilitate the elevated mitochondrial Ca^2+^ influx in AD HiNs (Fig. 2E), as increased MCU activity has been implicated in multiple AD models (3, 74, 75). The combined insults of excessive RyR-Ca^2+^ release with enhanced mitochondrial Ca^2+^ uptake and membrane depolarization (see below) trigger a large Ca^2+^ efflux from the mitochondria in AD HiNs (Fig. 2F), which may in part reflect increased mPTP opening (76–79). Notably, normalizing mitochondrial Ca^2+^ in AD neurons with Ryanodex suggests that the excessive ER-Ca^2+^ signaling is a dominant factor driving elevated mitochondrial Ca^2+^ levels.

### RyR-Ca^2+^ dynamics and mitochondrial membrane potential

Mitochondrial respiration and ATP production depend on a balance between Ca^2+^ influx and a highly negative membrane potential (36, 80). During activity-driven respiration, mitochondrial Ca^2+^ uptake causes a mild and transient depolarization of the membrane potential. This depolarization helps to prevent Ca^2+^ overload by reducing the proton motive force, which limits further Ca^2+^ influx. During these periods of elevated mitochondrial Ca^2+^, this membrane depolarization temporarily inhibits respiration and slows the flow of electrons along the ETC, thereby suppressing excessive ROS generation(81). This Ca^2+^-induced mild membrane depolarization is observed in RyR2 HEK cells (Fig 3H) and nonAD neurons (Fig 3K). However, AD neurons generate a markedly exaggerated depolarization of the ΔΨm in response to RyR-Ca^2+^ release, with membrane depolarization normalizing to nonAD levels after Ryanodex treatments (Fig. 2K). The disproportional depolarization in AD neurons is likely due to excessive mitochondrial Ca^2+^ uptake driven by dysregulated RyR-Ca^2+^ release (Fig 2C-E). Typically, mitochondrial Ca^2+^ efflux re-establishes the electrochemical gradient of the ΔΨm, allowing the mitochondria to resume normal respiration and ATP production. However, excessive mitochondrial Ca^2+^ influx and exaggerated depolarization of the ΔΨm in AD neurons during RyR-Ca^2+^ release may overwhelm the slower Ca^2+^ efflux mechanisms (Fig. 3F), therefore disrupting mitochondrial respiration. Prolonged disruption of the ETC can increase the risk of bioenergetic failure(82, 83), ROS production(84), and the release of pro-apoptotic factors(36, 85), all of which are implicated in AD pathology.

### Increased superoxide and caspase-3 generation in AD neurons

ROS are an inevitable byproduct of mitochondrial respiration, with mitochondrial Ca^2+^ influx increasing the activity of Ca^2+^-sensitive dehydrogenases (pyruvate, α-ketoglutarate, and isocitrate) within the TCA cycle and F_1_F_0_-ATP synthase complex (complex V) (13, 86, 87). Ca^2+^-enhanced production of NADH, the primary electron donor for the ETC, and the catalytic activity of F_1_F_0_-ATP synthase increase mitochondrial respiration. While physiological levels of Ca^2+^ are necessary to drive these essential metabolic processes, supraphysiological levels can cause an overproduction of NADH, which can excessively donate electrons to the ETC and promote electron leakage (18, 88). We observed increased levels of superoxides in AD neurons (Fig. 4A-B), which align with previous findings that have established ROS production and oxidative stress as common features of AD(63, 84). However, the pretreatment with Ryanodex reduced superoxide production in AD neurons to nonAD levels (Fig. 4-B), linking increased RyR-Ca^2+^ signaling as a driving factor for mitochondria-generated ROS in AD. In addition to increased mitochondrial Ca^2+^, the disrupted membrane potential in AD neurons (Fig. 3K) can impair ETC functionality, further increasing electron leakage and ROS production during RyR-Ca^2+^ signaling. These data provide a mechanistic connection between increased ER-mitochondrial Ca^2+^ signaling and ROS production in AD.

Under physiological conditions, mitochondrial Ca^2+^ influx (200 μM/s) is balanced by Ca^2+^ efflux (35-75 nM/s) via the NCLX (3, 89), and this homeostatic balance is calibrated to meet cellular demands for ATP synthesis. In AD neurons, the increased Ca^2+^ in the mitochondrial matrix drives exaggerated membrane depolarization (Fig. J-K) and increased ROS levels, all of which can promote mitochondria-induced apoptotic signaling via the intrinsic pathway (90–92). These AD-related pathologies are also linked to increased opening of the mitochondrial permeability transition pore (mPTP) (76). The permeabilization of the outer mitochondrial membrane through mPTP opening triggers a release of pro-apoptotic factors such as cytochrome c, which subsequently activates downstream caspases(93). This was evidenced in AD neurons by the detection of elevated caspase-3 activity (Fig. 4C), consistent with the increased mitochondrial Ca^2+^ load and its downstream effects on membrane depolarization and ROS production. However, 6-hour pretreatment with Ryanodex did not reduce caspase-3 activity, despite its ability to mitigate the aforementioned RyR-mediated mitochondrial dysfunctions (Fig. 4C). This suggests that the relatively short-term reduction in RyR-Ca^2+^ release in the AD neurons may not be sufficient to reverse the downstream caspase-3 cascade once it is activated. Beyond apoptosis, increased caspase-3 activity can induce DNA fragmentation, Aβ_42_ and tau tangle production via caspase-mediated cleavage, and heightened cellular stress, thus contributing on several levels to the progressive neurodegeneration associated with AD (94–98).

### Reduced Mitophagy in AD Neurons

Neurons rely on lysosome-mediated autophagy to remove cellular debris and maintain a homeostatic environment. Clearance of damaged organelles, including the mitochondria (via mitochondrial-autophagy or mitophagy), recycles damaged mitochondrial proteins and reduces reactive oxygen species (ROS) production and inflammation (48, 49). In this study, we demonstrated that mitophagy is impaired in the AD human neurons (Fig. 5A&B), likely due to defective lysosomal function resulting from Ca^2+^ dyshomeostasis (27). Specifically, chronically elevated Ca^2+^ levels disrupt vATPase proton pump activity and the trafficking of vATPase subunits to the lysosome, resulting in lysosomal alkalization and reduced lysosomal proteolytic function (27, 99, 100). To confirm that these disruptions are a product of lysosomal functional deficits and independent of lysosomal turnover, there are no differences in lysosome density between the nonAD and AD treatment groups (Supplemental Fig. 1B). Here, the partial rescue of mitophagy after normalization of RyR-Ca^2+^ signaling in AD HiNs indicates that aberrant Ca^2+^ signaling in AD human neurons prevents lysosomal-mediated removal of damaged mitochondria (Fig. 5B). Additionally, the accumulation of AD protein aggregates, like phosphorylated-tau, can prevent mitophagy initiator factors, like PARKIN, from being expressed on damaged mitochondrial membranes, thus reducing mitophagy and increasing oxidative stress (101). Furthermore, these findings suggest an inter-organelle dependency between the mitochondria and lysosomes, where both are disrupted by elevated ER-Ca^2+^ signaling in AD. In conjunction with this study, proteomic studies in both sporadic and familial AD brain tissues show differentially expressed levels of mitophagy-related signaling proteins, with increases in PINK1/Parkin, LC3-II and p62/SqSTM1 (102) and decreases in ATG5, ATG12, Beclin-1 (Bcl-1), AMBRA1, BNIP3, FUNDC1, VDAC1, and VCP/P97 (103). Both studies observed an accumulation of dysfunctional autophagosome vesicles with deficient mitophagic signaling. Several of these signaling molecules, including PINK1/Parkin, Bcl-1, MFN2, and VDAC, have been implicated at the ER-mitochondria interface, supporting the hypothesis that Ca^2+^ handling has a role in mitophagic regulation (49, 104, 105).

Here, we demonstrate key mechanisms by which dysregulated RyR-mediated Ca^2+^ release drives mitochondrial dysfunction and oxidative stress in AD. We continue to show that RyR regulation is crucial in restoring early AD pathology, such as proteinopathy, synaptic plasticity, calcium dysregulation, and now mitochondrial dysfunction. Attenuation of the RyR and decreasing the elevated cytosolic Ca^2+^ levels in human AD neurons fully rescued (or at least partially rescued), mitochondrial Ca^2+^ uptake, mitochondrial depolarization, superoxide production, caspase 3 production, and mitophagy. Thus, there is increasing compelling evidence demonstrating the early mechanisms by which disrupted intracellular Ca^2+^ handling through the RyR channel drives AD pathology, and highlights its potential as a therapeutic target for the treatment and prevention of AD.

## Materials and Methods

### Cell Culture

Stable, inducible human embryonic kidney (HEK293T) cells expressing wild-type (WT) RyR2 (RyR2 HEK) were obtained from Dr. Wayne Chen and are well characterized (106, 107). Cells were grown and maintained in Dulbecco’s modified Eagle’s medium (DMEM) supplemented with 0.1mM minimum Eagle’s medium nonessential amino acids, 4mM L-glutamine, 100IU/ml penicillin 100/mg of streptomycin, 4.5g glucose/liter, and 10% fetal calf serum, at 37°C under 5% CO_2_. Cells expressing the various RyR isoforms were selected using 200μg/ml hygromycin. Cells were plated on glass coverslips coated with 0.1% poly-L-lysine or 96-well Corning black, clear bottom plates, and given doxycycline to induce RyR expression 48 hours prior to use. Immunofluorescence was used to confirm the presence of RyR2 and RyR3. Cells in culture were treated with indicated concentrations of Ryanodex (10μM; 1–6 hours) and bafilomycin (25nM; 4 hours).

### iPSC and HiN Generation

Generation of induced pluripotent stem cell (iPSC)-derived human induced neurons (HiN) has been previously described (108). Briefly, normal control (nonAD) and familial-AD patient fibroblasts (A246E PS1 mutation) obtained from the Coriell Institute were converted to iPSCs using gene delivery of the Yamanaka factors by vector or RNA transfection (ReproRNA-OKSGM kit including five vectors OCT4, KLF-4, SOX2, GLIS1, and c-MYC). Fibroblasts were transfected and cultured for 14–21 days in REPRO-TeSR medium. Viable iPSC colonies were converted to HiN by lentiviral vectors containing the transcription factor NGN2. HiNs were selected for lentiviral transduction by puromycin resistance, allowed to mature over 14–28 days in neural basal media (ThermoFisher) with doxycycline, brain-derived neurotrophic factor, and neurotrophin 3 (PeproTech). Induced neurons were fed with 50:50 mixture of astrocyte-conditioned medium (provided by Dr. Allison Ebert). HiN were grown on Matrigel-coated coverslips or 24-well (Costar) plates. Cells were incubated with indicated concentrations of vehicle, Ryanodex (10μM; 1-6 hours), or bafilomycin A1 (25nM; 4 hours) before running physiological experiments.

### Mitochondrial Calcium Imaging

Cells were incubated with crude lysate of AAV9-hSyn-2mt-Mito-GCaMP6m (0.3ul/well, courtesy of Dr. Israel Sekler (109, 110) for 3 days (RyR2 HEK cells) or 7 days (human iPSC-derived neurons). This is a modified Ca^2+^ indicator protein variant, mitoGCaMP6m, containing the neuron-specific promoter (hSyn=synapsin) and a double mitochondrial targeting sequence (2mt) that is delivered using adeno-associated viral vector AAV2/9. Cells were imaged on a modified upright Olympus BX51WI microscope. Cells were continuously perfused with oxygenated (95% O_2_-5% CO_2_) artificial cerebrospinal fluid (aCSF) [125mM NaCl, 2.5 KCl, 1.25mM KH_2_PO_4_, 10mM dextrose, 25mM NaHCO_3_, 2mM CaCl_2_, and 1.2mM MgSO_4_, pH 7.4] at room temperature. The microscope is coupled to a Lumencor Spectra X light source. The fluorescence responses were captured using a 40X water-immersion Olympus objective and analyzed using Nikon NIS-Elements Software (AR package). Bath application of caffeine (10mM, Sigma) was used to evoke RyR-evoked calcium release. Bath application of CCCP (5μM, Sigma) was used as a positive control for mitochondrial depolarization. Ryanodex (10μM; 1 hour; Lyotropic Therapeutics Inc.) was dissolved in aCSF and used as a negative allosteric modulator of RyR. Mitochondrial Ca^2+^ is reported as a percentage of the peak response over average baseline after background subtraction. Mitochondrial Ca^2+^ influx and efflux was calculated by a linear fit of the change in the fluorescence before and after the peak fluorescence, as described in (110).

### Mitochondrial Membrane Potential Imaging

Cells were incubated with either Rhod123 (ThermoFisher, 10μM, 30min incubation followed by 15min calibration in 0.1M PBS at at 37°C under 5% CO_2_) or tetramethylrhodamine methyl ester (TMRM, ThermoFisher, 50nM, 1hr). Cells were washed in 0.1M PBS then imaged on a modified upright Olympus BX51WI microscope. The fluorescence responses were captured using a 40x water-immersion Olympus objective and analyzed using Nikon NIS-Elements Software (AR Package). Cells were continuously perfused with oxygenated (95% O_2_, 5% CO_2_) artificial cerebrospinal fluid (aCSF) [see above for more details] at room temperature. The microscope is coupled to a Lumencor Spectra X light source. Bath application of caffeine (10mM, Sigma) was used to evoke RyR-evoked calcium release, followed by a perfusion of CCCP (5 μM; Sigma) as positive control for mitochondrial depolarization. Ryanodex (10μM; 1 hour; Lyotropic Therapeutics Inc.) was dissolved in aCSF and used as a negative allosteric modulator of RyR. After background subtraction, resting (TMRM) mitochondrial membrane potential was calibrated to CCCP-generated maximum fluorescent (F/Fmax), while RyR-driven depolarization was reported as percentage of the peak response over average baseline.

### Mitochondrial Superoxide Imaging

To measure mitochondrial superoxide levels, human-induced neurons were incubated with Ryanodex (10μM, 1 hour) with MitoSOX Deep Red dye for 30 min at 37°C with 5% CO_2_ (0.1μM; Dojindo Molecular Technologies, Inc) and imaged at the far-red fluorescence spectrum (λex: 540 nm, λem: 670 nm). A 40X water-immersion objective using an upright Olympus BX51WI microscope with a Lumencor Spectra X light source. Images were analyzed on ImageJ (FIJI) and presented as mean fluorescent intensity above background. In cell culture, cell bodies from brightfield images were counted using the FIJI Cell Counter plug-in. No differences were seen in the average number of cells per treatment group (Supplemental Fig. 1A)

### Caspase 3/7 Activity Assay

Caspase 3 activity was measured using commercially available Caspase-Glo 3/7 Assay (G809; Promega) following the manufacturer’s instructions for a 24-well plate. Ryanodex-treated neurons were incubated in Ryanodex (10μM; 6 hours), while control groups were incubated with DMSO (0.1%). The luminescent signal, after background subtractions, was measured (λ: 510nm) using a FlexStation 3 Automated Cell Plate Reader acquiring a 3-point well-scan recording.

### Mitophagy Imaging

Following treatments, neurons were incubated with mitophagy dye at 37°C with 5% CO_2_ (0.1μM; 30 min.; Dojindo Molecular Technologies, Inc; Mitophagy Detection Kit), then incubated with lysosome dye at 37°C with 5% CO_2_ (1μM; 30 min.; Dojindo Molecular Technologies, Inc). Neurons were then imaged with a 40X water-immersion objective using an upright Olympus BX51WI microscope with a Lumencor Spectra X light source. Images were analyzed on Metamorph software (version 7) and presented as mean fluorescent density above background or analyzed on FIJI and presented as mean fluorescence intensity above background, where appropriate. In cell culture, cell bodies from brightfield images or DAPI staining were counted using the FIJI Cell Counter plug-in. The fluorescence density was standardized to cell count. No differences were seen in the average number of cells per treatment group (Supplemental Fig. 1B)

### Experimental Design and Statistical analyses

Statistical analysis was performed using Graph Pad Prism 7. Data are represented as mean ± SE. Unpaired t-tests, one-way ANOVA or two-way ANOVA, with Tukey’s post hoc analysis, were performed where appropriate. Statistical significance was set at p<0.05.

## Acknowledgments

Dr. Allison Ebert for graciously providing Astrocyte Conditioned Media for the generation of human-induced neurons. Dr. Wayne Chen for graciously providing the RyR2-expressing HEK293T cell line.

## Figure Legends

**Supplemental Figure 1.**
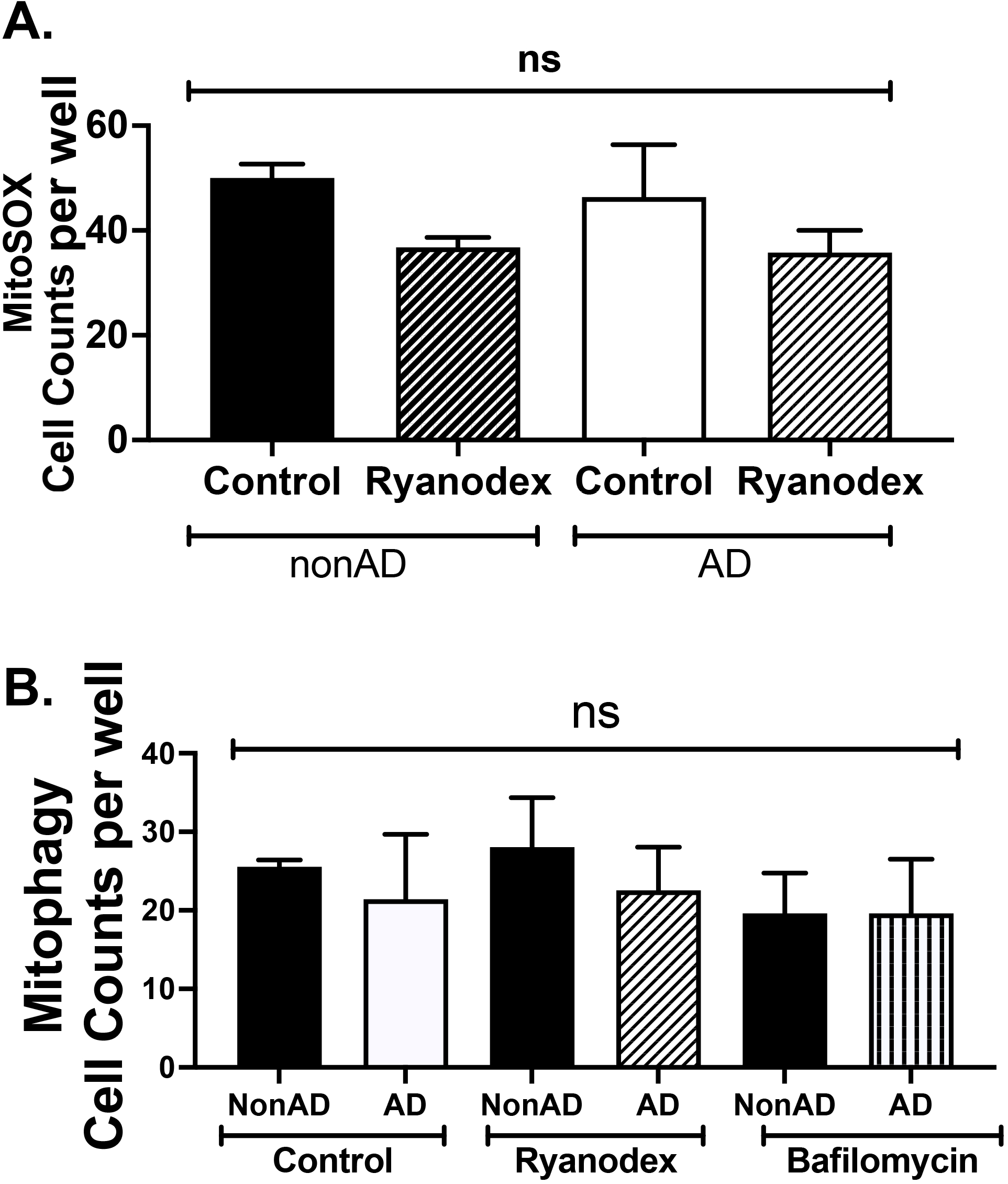
(A) For the mitochondrial superoxide study, cell counts showed no difference in number of human neurons amongst treatment groups for both nonAD and AD. (B) For the mitophagy study, cell counts showed no difference in number of human neurons amongst treatment groups for both nonAD and AD.

## Notes

### Competing Interest Statement

The authors have declared no competing interest.

